# MANGEM: a web app for Multimodal Analysis of Neuronal Gene expression, Electrophysiology and Morphology

**DOI:** 10.1101/2023.04.03.535322

**Authors:** Robert Hermod Olson, Noah Cohen Kalafut, Daifeng Wang

## Abstract

Single-cell techniques have enabled the acquisition of multi-modal data, particularly for neurons, to characterize cellular functions. Patch-seq, for example, combines patch-clamp recording, cell imaging, and single-cell RNA-seq to obtain electrophysiology, morphology, and gene expression data from a single neuron. While these multi-modal data offer potential insights into neuronal functions, they can be heterogeneous and noisy. To address this, machine-learning methods have been used to align cells from different modalities onto a low-dimensional latent space, revealing multi-modal cell clusters. However, the use of those methods can be challenging for biologists and neuroscientists without computational expertise and also requires suitable computing infrastructure for computationally expensive methods. To address these issues, we developed a cloud-based web application, MANGEM (Multimodal Analysis of Neuronal Gene expression, Electrophysiology, and Morphology) at https://ctc.waisman.wisc.edu/mangem. MANGEM provides a step-by-step accessible and user-friendly interface to machine-learning alignment methods of neuronal multi-modal data while enabling real-time visualization of characteristics of raw and aligned cells. It can be run asynchronously for large-scale data alignment, provides users with various downstream analyses of aligned cells and visualizes the analytic results such as identifying multi-modal cell clusters of cells and detecting correlated genes with electrophysiological and morphological features. We demonstrated the usage of MANGEM by aligning Patch-seq multimodal data of neuronal cells in the mouse visual cortex.

**Author Summary:** The human brain is made up of billions of tiny cells called neurons, each with their own important job. Scientists are now able to study individual neurons in more detail than ever before using new advanced techniques. They can look at different data of individual neurons like how genes are being used (gene expression), how the neuron responds to electrical signals (electrophysiology), and what it looks like (morphology). By combining all of this information, they can start to group similar neurons together and figure out what they do. However, due to the data complexity, this process can be very complicated and hard for researchers without sufficient computational skills. To address this, we developed a web app, MANGEM (Multimodal Analysis of Neuronal Gene Expression, Electrophysiology, and Morphology). It lets scientists upload their data and select emerging machine-learning approaches to find groups of similar neurons. It also makes interactive visualizations to help them explore the characteristics of neuron groups and understand what they do.

## Introduction

The human brain has approximately 86 billion neurons encompassing a vast range of different functions. Understanding the roles of individual neurons is a daunting challenge that is beginning to become possible with new techniques and technologies. The development of single-cell technologies such as Patch-seq has resulted in the ability to characterize neurons with new specificity and detail. Patch-seq enables a researcher to simultaneously obtain measures of gene expression, electrophysiology, and morphology of individual neurons. Gene expression is a measure of the extent to which different genes in a cell’s DNA are transcribed to RNA and then translated to produce proteins. Electrophysiology describes the electrical behavior of a cell. A microscopic pipette containing an electrolyte contacts the cell membrane to establish an electrical connection. Then the cell’s electrical response to an applied voltage or current is measured. Morphology refers to the physical structure of a neuron, including the size and shape of the cell’s axon and dendrites. By combining microscopy, RNA sequencing, and electrophysiological recording for individual neurons, multi-modal datasets can be developed with the potential to reveal relationships between neuronal function, structure, and gene expression (1). Multi-modal single-cell datasets are increasingly available to researchers, in part due to efforts by the Brain Research through Advancing Innovative Neurotechnologies (BRAIN) Initiative to support the development and storage of such datasets in freely accessible repositories such as the Neuroscience Multi-Omic Archive (2) for genomic data and Distributed Archives for Neurophysiology Data Integration (3,4) for neurophysiology data, including electrophysiology.

While multi-modal single-cell data offers great potential for improving understanding of brain organization and function, new methods are required for integration and analysis of the data (5). Because cells with similar characteristics in one modality are not necessarily similar when measured by another, identification of cell clusters must incorporate disparate data types simultaneously. Machine-learning methods such as manifold learning are highly applicable to the problems posed by heterogeneity of multi-modal single-cell data (6), but these methods are commonly difficult to use, especially for biologists and neurologists who may not have computational expertise. Documentation and tutorials, if present, are limited in scope. The methods are often supplied as source code only, requiring coding expertise to use, which further limits their accessibility. Installation and configuration of the software adds another layer of difficulty to overcome before these methods can be applied. As an example, consider the software for UnionCom (7). While the UnionCom software is available in the Python package index (PyPI) and easily installable, its dependencies are not automatically installed. The prospective user will quickly discover that the versions of those dependencies suggested in the limited documentation are not easily installable in recent versions of Python. Given time and effort, a motivated researcher will manage to find the right combination of package versions and Python version that will be compatible, but this level of difficulty is both a significant barrier to use and common in open-source scientific software generally (8).

An increasingly common way to address the challenges of running open-source scientific software is by implementing the methods of the software in a web application (9,10). Here we present a new web application named MANGEM (Multimodal Analysis of Neuronal Gene Expression, Electrophysiology and Morphology), developed to address the challenges researchers may experience in using existing methods of aligning and analyzing multi-modal single-cell data. In particular, MANGEM provides an easy-to-use interface to a variety of machine-learning alignment methods, requires no coding to use, and does not require installation of software or management of computing infrastructure. Preloaded datasets and an interface that walks the user through each operational step provide for an accessible introduction to the use of machine-learning methods to align multi-modal datasets. As a cloud-based web application, MANGEM enables users to begin exploring multi-modal single-cell datasets without first undertaking the challenges of software installation or management of the underlying infrastructure. While the application was designed for real-time data processing and exploration, it also supports running certain long-running methods asynchronously, providing a customized URL for users to retrieve results after computation is complete. Interactive graphical display of output facilitates exploration of the data at each step of the analysis process: raw data as uploaded, preprocessed data (e.g., standardized), aligned datasets, and cross-modal clusters. Integrated downstream analysis methods support identification of important cellular features within cross-modal cell clusters and aid interpretation of the revealed relationships within cell clusters.

## Design and Implementation

The MANGEM web application offers a range of methods for aligning multi-modal data of neuronal cells, identifying cross-modal cell clusters using the aligned data, and generating visualizations to facilitate the characterization of these cross-modal clusters, including their differentially expressed genes and correlated multi-modal features (**Fig. 1**).

**Figure 1.**
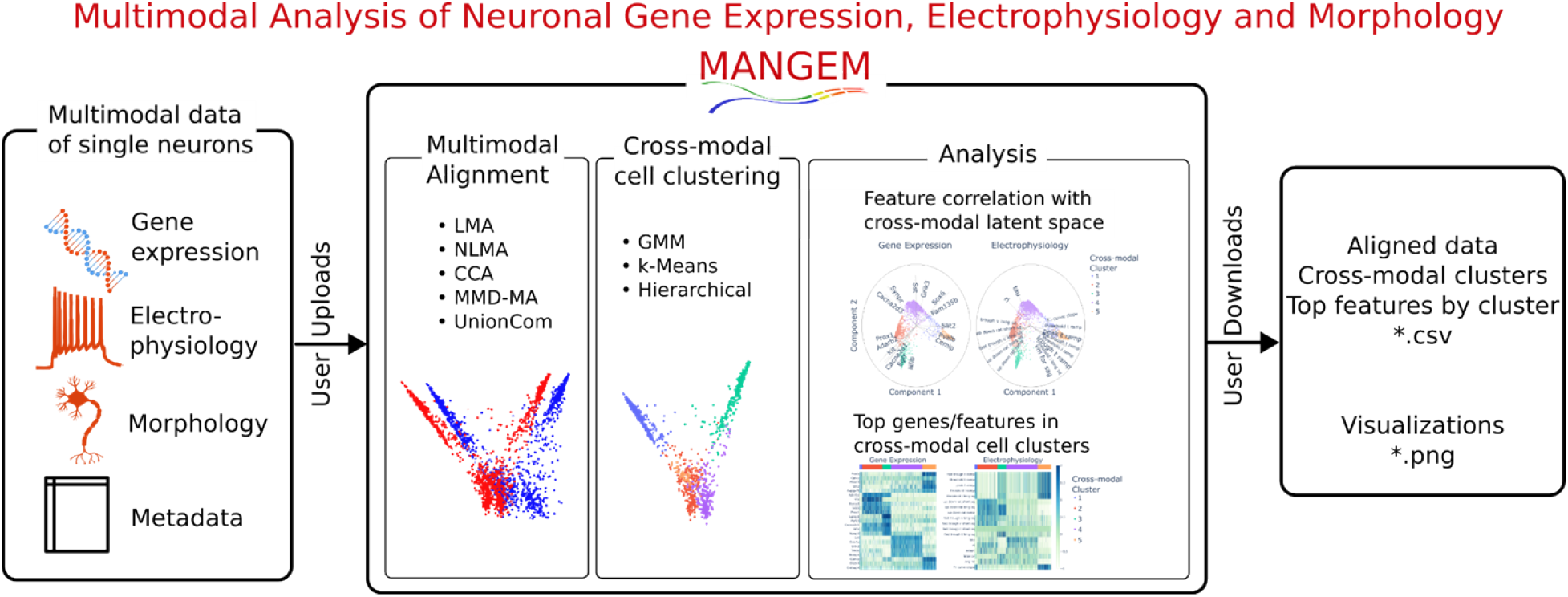
Overview of MANGEM (Multimodal Analysis of Neuronal Gene Expression, Electrophysiology and Morphology). User input to MANGEM includes multimodal single-cell data together with cell metadata. Within MANGEM, the multimodal data are aligned using machine learning methods, projecting disparate modalities into a low-dimensional common latent space. Clustering algorithms are applied within the latent space to identify cell clusters, and then analysis methods are provided in MANGEM to characterize the clusters by differential feature expression and correlation of features with the latent space. In addition to interactive plots generated at each step of the workflow, downloadable output includes tabular data files (cell coordinates in latent space, cluster annotations, top features for each cluster) and images depicting alignment, cross-modal cell clusters, and cluster analyses.

The application is implemented using Plotly Dash Open Source, a Python-based framework for developing data science applications (11). Dash is based on Plotly.js (12), React (13), and Flask (14), and it functions by tying user interface elements to stateless callback functions. In the case of MANGEM, some callback functions are quasi-stateless, in that uploaded and aligned datasets are stored in a filesystem cache to avoid repeating lengthy calculations.

Our public deployment of MANGEM is on Amazon Web Services infrastructure (**Fig. 2**). The Elastic Beanstalk service is used to deploy the application to an Elastic Compute Cloud (EC2) instance with associated storage in Amazon Simple Storage Service (S3). In order to be accessed by a user with a web browser, MANGEM requires additional software. A reverse proxy server directs the requests from the web browser to an application server that can translate the requests to the Web Server Gateway Interface (WSGI) protocol used for communication with MANGEM. By default, the Elastic Beanstalk Python platform provides nginx (15) as the reverse proxy server and Gunicorn (16) as the WSGI application server; however, MANGEM does not depend on those specific programs. For example, in our development environment, we use the Apache HTTP server with mod_proxy (17) as the reverse proxy server and uWSGI (18) as the WSGI application server. Most data processing occurs within the main MANGEM process, but additional software is required to enable long-running alignment jobs to run asynchronously. In this case, Celery (19) is used to run those background jobs, and Redis (20) is used as a message broker to communicate between MANGEM and Celery. Whether aligned synchronously or asynchronously, aligned multi-modal datasets are stored in a filesystem cache on AWS S3.

**Figure 2.**
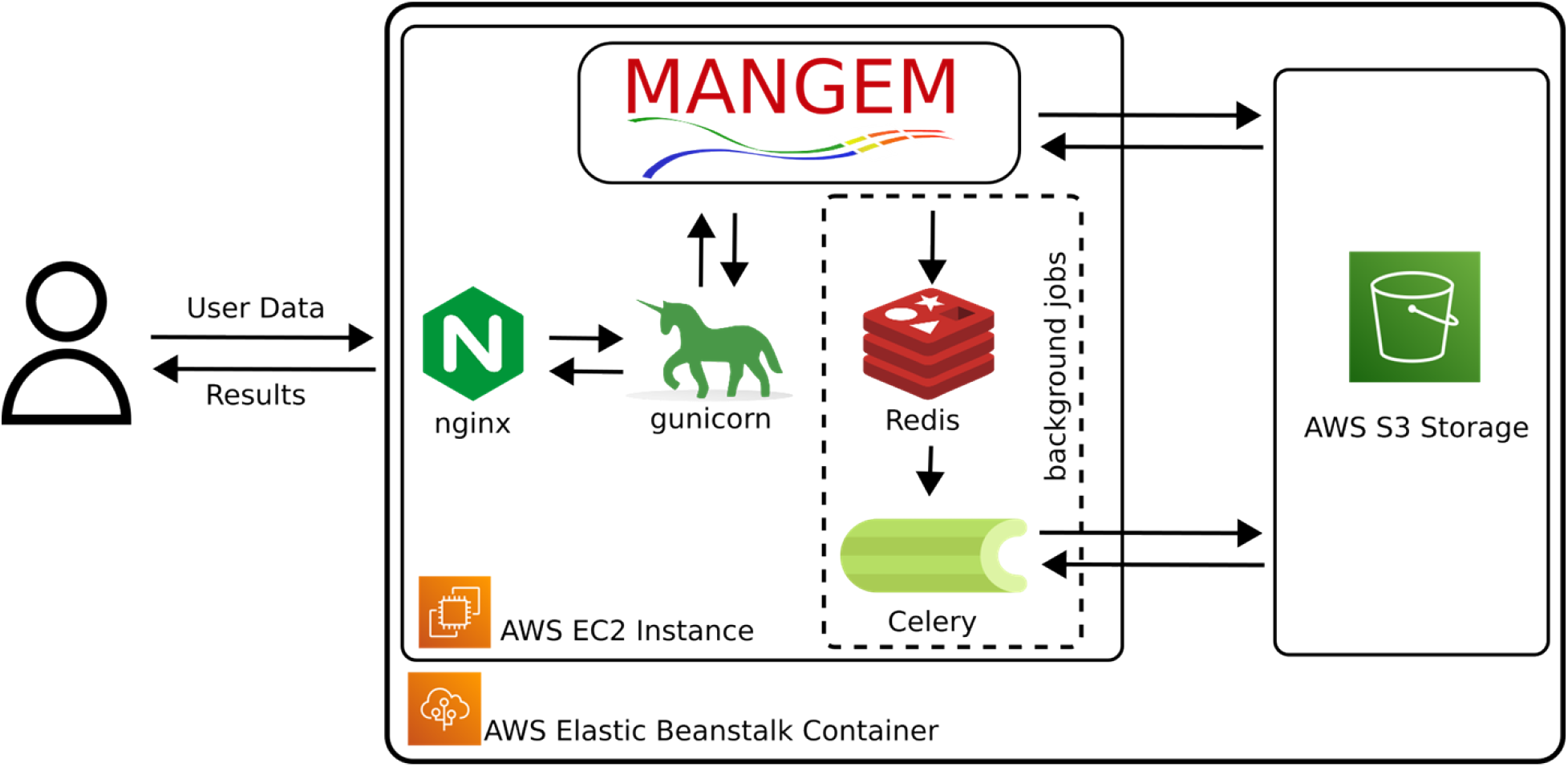
Cloud implementation of MANGEM using AWS infrastructure. The application runs on Amazon Cloud Services using Elastic Beanstalk to provision an EC2 instance. The web server nginx serves as a reverse proxy to the gunicorn WSGI server. MANGEM is written in Python using the Plotly Dash framework. Long-running tasks are run in the background by Celery workers, with Redis acting as the message broker between MANGEM and Celery. Uploaded and processed data files are stored in a filesystem cache in AWS S3.

MANGEM’s layout is organized as a set of tabs on the left which contain user interface controls, while the right side contains plots or other information related to the active tab. The tabs correspond to the sequence of steps users will typically take when running the application: upload data, align data, identify cross-modal cell clusters, and perform downstream analysis of cross-modal cell clusters. Each tab contains controls that allow the user to adjust parameters relevant to the current step of the workflow and which influence the downstream results (**Table 1**).

**Table 1.** Key parameters of data processing and analysis in MANGEM. The listed parameters all influence downstream output of MANGEM. For example, selecting a preprocessing method on the Upload Data tab will result in that method being applied to the uploaded dataset before the selected multi-modal alignment method is applied.

| MANGEM functions | Parameter |
| --- | --- |
| Step 1. Upload Data | Preprocessing method for each modality (log/standardize) |
| Step 2. Alignment | Alignment algorithm<br>Dimension of machine learning latent space (3-10)<br>Number of nearest neighbors (1-10) used in constructing |
|  | similarity matrices (LMA, NLMA).<br>Number of iterations (MMD-MA) |
| Step 3. Clustering | Clustering algorithm<br>Number of cell clusters (1-10) |
| Step 4. Analysis | Component selection<br>Number of top features per cell cluster in Features of Cross-modal Clusters<br>Number of top correlated features in Top Feature Correlation with Latent Space |

At each step of the workflow (**Fig. 3**), interactive figures are automatically generated to support understanding, and computation products are available for download as tabular data files. A video demonstration of the workflow is provided in the Supplemental Data.

**Figure 3.**
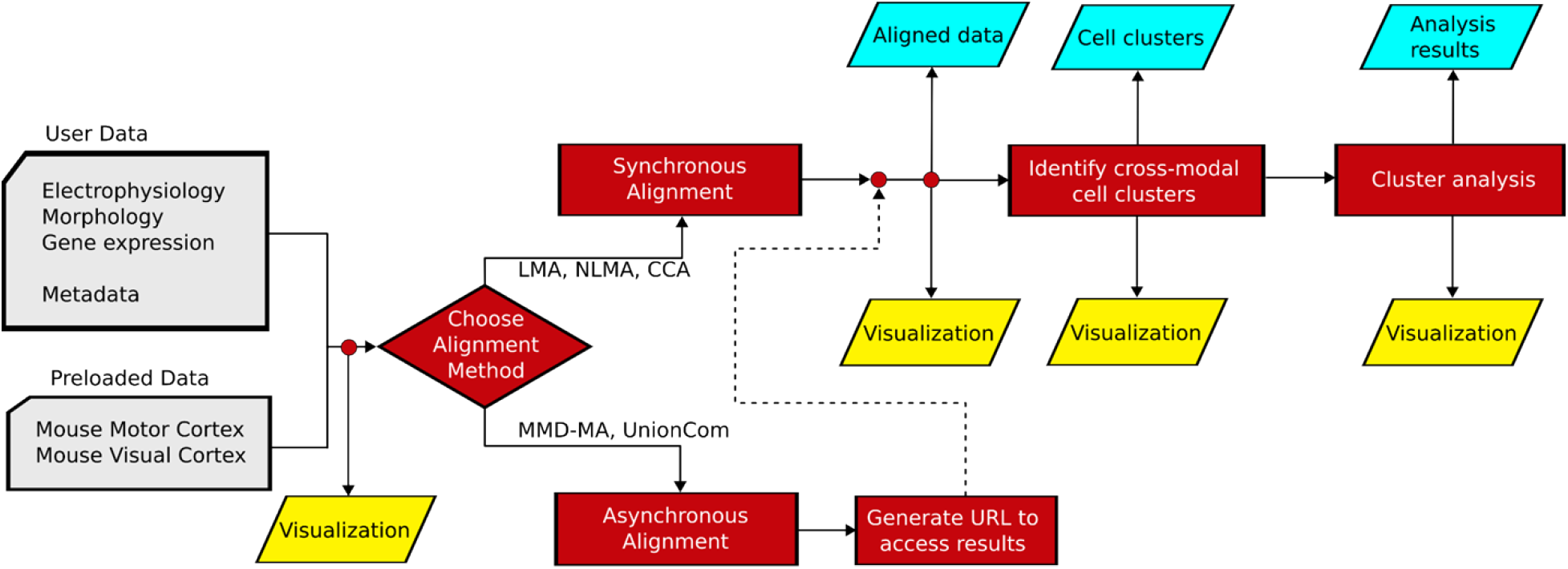
Data flow through MANGEM web application. Input data passes into an alignment process, which will either run in the main process or in the background, depending on the method. In the case of background (asynchronous) alignment, a URL will be supplied to the user which will allow them to check on the job’s status and access the results upon completion. Aligned data feed into a clustering algorithm, and then data analysis methods can be applied to the cell clusters. Data visualization output can be produced at each stage of the process, and tabular data files of aligned data, cell clusters, and analysis results can be downloaded.

### Step 1: Upload Data

The first data processing step in MANGEM is selecting or uploading neuronal data, accomplished on the Upload Data tab. The expected input to MANGEM consists of three data files in .csv format: one file for each of two modalities, and a third file of cellular metadata. Three sample data sets are preloaded in MANGEM, and a link is provided within the application to download one of these. The first column of each file should contain a cell identifier, and the files are expected to have a consistent cell order. Denote data for the first modality as *X*, data for the second modality, *Y*, and metadata, *M*. Each of these has *n* rows corresponding to *n* neurons. *X* and *Y* have *d*_1_ and *d*_2_ features, respectively. Metadata matrix *M* has *d_m_* cell characteristics.

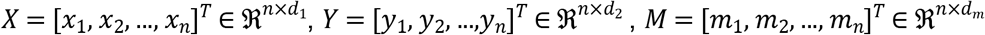

When user data is uploaded, a label may be supplied for each modality; otherwise, the modalities will be identified using the default labels of “Modality 1” and “Modality 2” in plot legends.

A preprocessing operation may optionally be selected for each modality. Choices include “Log transform” and “Standardize”, which for the first modality would be:

“Log transform”: 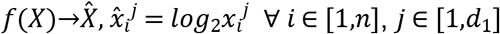
“Standardize”: 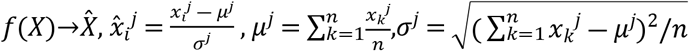

If a preprocessing operation is selected, that operation will be applied to the appropriate dataset prior to alignment. The default values of “Log transform” for Modality 1 and “Standardize” for Modality 2 are suitable for the preloaded datasets, where Modality 1 is Gene Expression and Modality 2 is Electrophysiology.

### Data Exploration

The “Explore Data” section of the Upload Data tab can generate plots to gain insight into cell features in the uploaded or selected data sets. A series of box plots is generated for each value of a categorical metadata variable when a single cell feature is selected (**Fig. S1a**). A particular value of that metadata variable may be selected to filter the data, in which case a violin plot is generated (**Fig. S1b**). It is also possible to select two features to compare in a scatter plot (**Fig. S1c**). These features could be from the same or different modalities. Similarly, to the single-feature case, selecting a specific value of a metadata variable filters the data so that only the points corresponding to cells having that metadata value are displayed in the scatter plot (**Fig. S1d**). As with all plots in MANGEM, a toolbar will pop up when the cursor is placed over the plot. The toolbar has buttons to change to plot appearance (zoom or pan, for example) and also has a button with a camera icon that causes an image of the plot to be downloaded.

### Step 2: Multi-modal Alignment

The approach used by MANGEM to find clusters of related cells is to first transform the measured cellular features into a latent space where cells having similar features are closer together. This transformation process is called multi-modal alignment, and several alignment methods are implemented in MANGEM. Currently supported methods include Linear Manifold Alignment (LMA), Nonlinear Manifold Alignment (NLMA) (21), Canonical Correlation Analysis (CCA), Manifold Alignment with Maximum Mean Discrepancy (MMD-MA) (22), and Unsupervised Topological Alignment for Single-Cell Multi-Omics (UnionCom) (7). LMA and NLMA utilize similarity matrices to formulate a common latent space. MMD-MA minimizes an objective function which measures distortion and preserved representation. UnionCom infers cross-modal correspondence information before using t-SNE (23) to provide the final latent spaces.

Several parameters of these alignment methods can be adjusted on MANGEM’s Alignment tab. These include the dimension of the latent space, the number of nearest neighbors to be used when computing the similarity matrix (LMA, NLMA), and the number of iterations (MMD-MA). The alignment methods take as input the preprocessed datasets 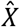 and 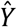 if no preprocessing method has been selected, then data are used as uploaded: 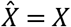 and 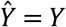. If we think of the alignment as finding optimal projection functions *f* and *g* which project cellular data from modality 1 and modality 2, respectively, to a common latent space of dimension *d*, then after alignment, the *i^th^* cell can be represented by 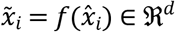 and 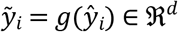.

After alignment has been completed, the cellular coordinates in the latent space can be downloaded by clicking on the “Download Aligned Data” button on the Alignment tab of MANGEM.

### Asynchronous Computation of Alignment

Though MANGEM primarily operates synchronously, some of the supported alignment methods (notably, UnionCom and MMD-MA) require enough computational resources to motivate running those tasks in the background, asynchronously. Celery, an open-source asynchronous task queue, is used to queue and run these long-running alignment tasks in the background. When the user clicks the “Align Datasets” button after selecting the UnionCom or MMD-MA alignment method, the alignment job is submitted to the task queue, and a unique URL is provided to the user. Navigating to this URL will give the user a message indicating the job status: waiting to start in the task queue, running, or complete. If the job is complete, then the results will be loaded and the Clustering tab of MANGEM will open, and the usual clustering and analysis methods will be available. A video demonstration of background alignment is included in Supplemental Data.

### Step 3: Cross-modal Cell Clustering

Once the multi-modal single-cell data have been aligned, cell clusters can be identified based on proximity within the latent space. Three different clustering methods are currently supported by MANGEM: Gaussian mixture model, K-means, and hierarchical clustering, all using methods provided by the Scikit-learn Python package (24). Gaussian mixture model clustering uses the GaussianMixture class with a single covariance matrix shared by all components and 50 iterations. K-means uses the KMeans class with the parameter n_init set to 4 and random seed specified. Hierarchical clustering is implemented using the AgglomerativeClustering class with Ward linkage, which minimizes the sum of squared distances within clusters. In all cases, the number of clusters to be identified can be specified using the slider control on the Clustering tab of MANGEM. After clusters have been identified, the assignment of cells to clusters can be downloaded by clicking on the “Download Clusters” button.

### Step 4: Analysis of Cross-modal Cell Clusters

The Analysis tab of MANGEM supports visualization of alignment and clustering results as well as methods to reveal relationships between cell features in the context of identified cell clusters. These methods are accessed via the Plot type selection control.

### Features of cross-modal clusters

The “Features of cross-modal clusters” method identifies the most important features within each cross-modal cluster and generates a heatmap for each modality where the rows correspond to identified features and the columns correspond to cells, grouped into previously identified clusters. The number of features identified for each cluster is specified using the “Number of Top Features per cluster” control on the Analysis tab. A list of the most important features can be downloaded using the “Download Top Features” button.

### Top feature correlation with latent space

Top feature correlation with latent space creates a bibiplot (i.e., collection of biplots) (25) with one biplot for each modality. Each biplot displays a 2-dimensional projection of the aligned data for the modality in the latent space while overlaying lines corresponding to the features which are most highly correlated with the latent space representation. For each modality, the correlation is computed between the original cellular data for each feature and the projection of the cellular data into the latent space dimensions selected as components X and Y on the Analysis tab. For a given feature, the correlations between that feature and its X and Y latent space representation determine the coordinates of the endpoint of that feature’s line.

The latent space dimensions in which aligned data are plotted can be selected using the “Component Selection” controls on the Analysis tab. At most, three dimensions can be plotted at one time within MANGEM, but these controls allow the user to select which dimensions are plotted to gain different perspectives on the data. Additional controls on the Analysis tab allow aligned data points to be colored either by cluster or by metadata value (for example, transcriptomic cell type).

## Results

In this study, we showcased the usage of MANGEM through two case studies that utilized emerging Patch-seq multimodal data of inhibitory neuronal cells in the mouse visual cortex (such as gene expression, electrophysiology, and morphology). It is worth noting that MANGEM is a general-purpose tool that can be used for any user multimodal data of neurons.

### Case Study 1: neuronal gene expression and electrophysiology

We first tested MANGEM to align these neuronal cells based on gene expression and electrophysiological features. We uploaded two datasets, one containing 1302 most variable expressed genes and 41 electrophysiological features for 3654 neuronal cells, on the Upload Data tab of MANGEM. We then preprocessed the data using log transformation for gene expression and standardization for electrophysiology features.

On the Alignment tab, we set the alignment method to Nonlinear Manifold Alignment (NLMA), the number of latent space dimensions to 5, and the number of nearest neighbors (used in construction of the similarity matrix for NLMA) to 2. Clicking the “Align Datasets” button generated two measures of alignment along with a 3D plot of the aligned cells (**Fig. 4a**). The aligned multimodal cells were represented in the common latent space, 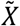 and 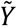, are 3654 cells (rows) by 5 latent dimensions (columns). We also tested other alignment methods and found that NLMA, in addition to being one of the fastest methods to run, resulted in the smallest alignment error (**Fig. S2, Table S1**).

**Figure 4.**
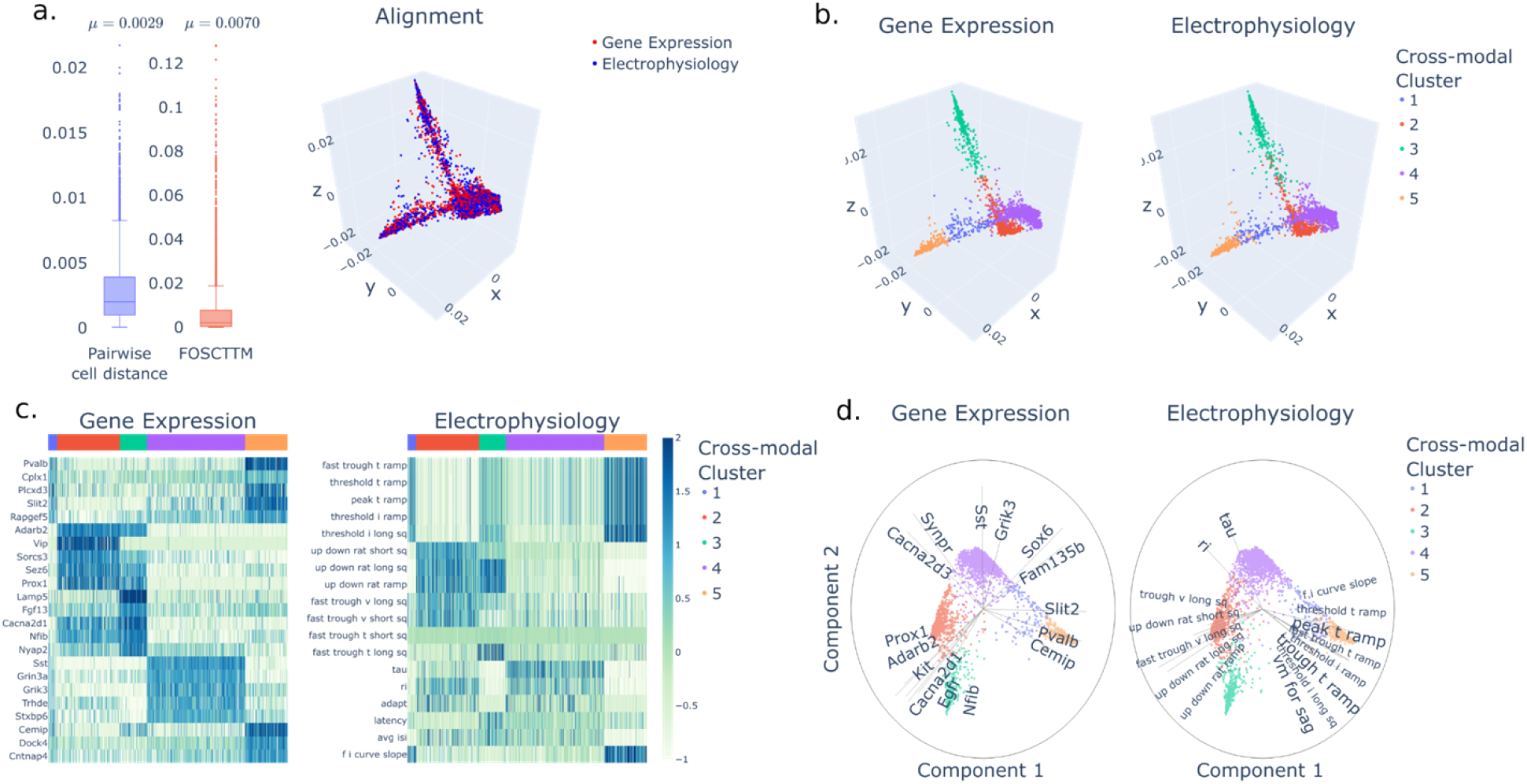
MANGEM analysis and visualization of neuronal gene expression and electrophysiological features in mouse visual cortex. a.) Measures of alignment error and 3-d plot of superimposed aligned data in latent space are shown for the preloaded mouse visual cortex dataset after nonlinear manifold alignment. b.) Cross-modal clusters, obtained by Gaussian mixture model, are indicated by color in plots of aligned data for each modality. c.) Feature levels across all cells for the top 5 features for each cross-modal cluster. Normalized feature magnitude was ranked using the Wilcox Rank Sum test. Cross-modal clusters are identified by the colored bar at the top of each plot. d.) Biplots for Gene Expression and Electrophysiological features using dimensions 1 and 2 of the latent space. The top 15 features by correlation with the latent space are shown plotted as radial lines where the length is the value of correlation (max value 1).

Afterwards, we chose to use the Gaussian Mixture Model clustering algorithm on the Clustering tab, specifying 5 clusters. Upon clicking the “Identify Cross-Modal Cell Clusters” button, the algorithm identified cross-modal cell clusters and generated side-by-side plots of the aligned cells for each modality in the latent space. These plots showed cells colored according to their respective cross-modal clusters (**Fig. 4b**).

The Analysis tab of MANGEM offers various visualization methods for exploring cross-modal relationships between gene expression and electrophysiological features. We set the number of top features to 5 and selected the “Features of Cross-modal Clusters (Heatmap)” (**Fig. 4c**). The resulting heatmap showed that tau and ri were the top two electrophysiological features in Cluster 4, while the top differentially-expressed genes in the cluster were Sst, Grin3a, Grik3, Trhde, and Stxbp6. These shared multi-modal features suggest potential functional linkages among the cells in the cluster.

To further investigate these linkages, we switched the plot type to “Top Feature Correlation with Latent Space (Bibiplot)” (**Fig. 4d**) and set the number of top correlated features to 15. The bibiplots graphically represented the most highly-correlated features from cross-modal cell clusters and allowed for interactive zooming into the Cluster 4 area on the latent space. The highly-correlated features included tau and, to a lesser extent, ri among the electrophysiological features, while Sst and Grik3 were among the genes associated with Cluster 4.

### Case Study 2: neuronal gene expression and morphology

MANGEM was used to process gene expression of the top 1000 variable genes and morphological features in the mouse motor cortex (25). The data consists of 646 single-cells with 42,466 genes and 63 morphological features. Each modality is formatted into a separate csv, with an additional file indicating metadata such as age, gender, etc. The data was then uploaded onto the webapp using the upload tab.

MANGEM can be used to easily test multiple integration methods. For this application, we chose Non-Linear Manifold Alignment (NLMA). After alignment, pairwise accuracy statistics are reported (**Fig. 5a**).

**Figure 5.**
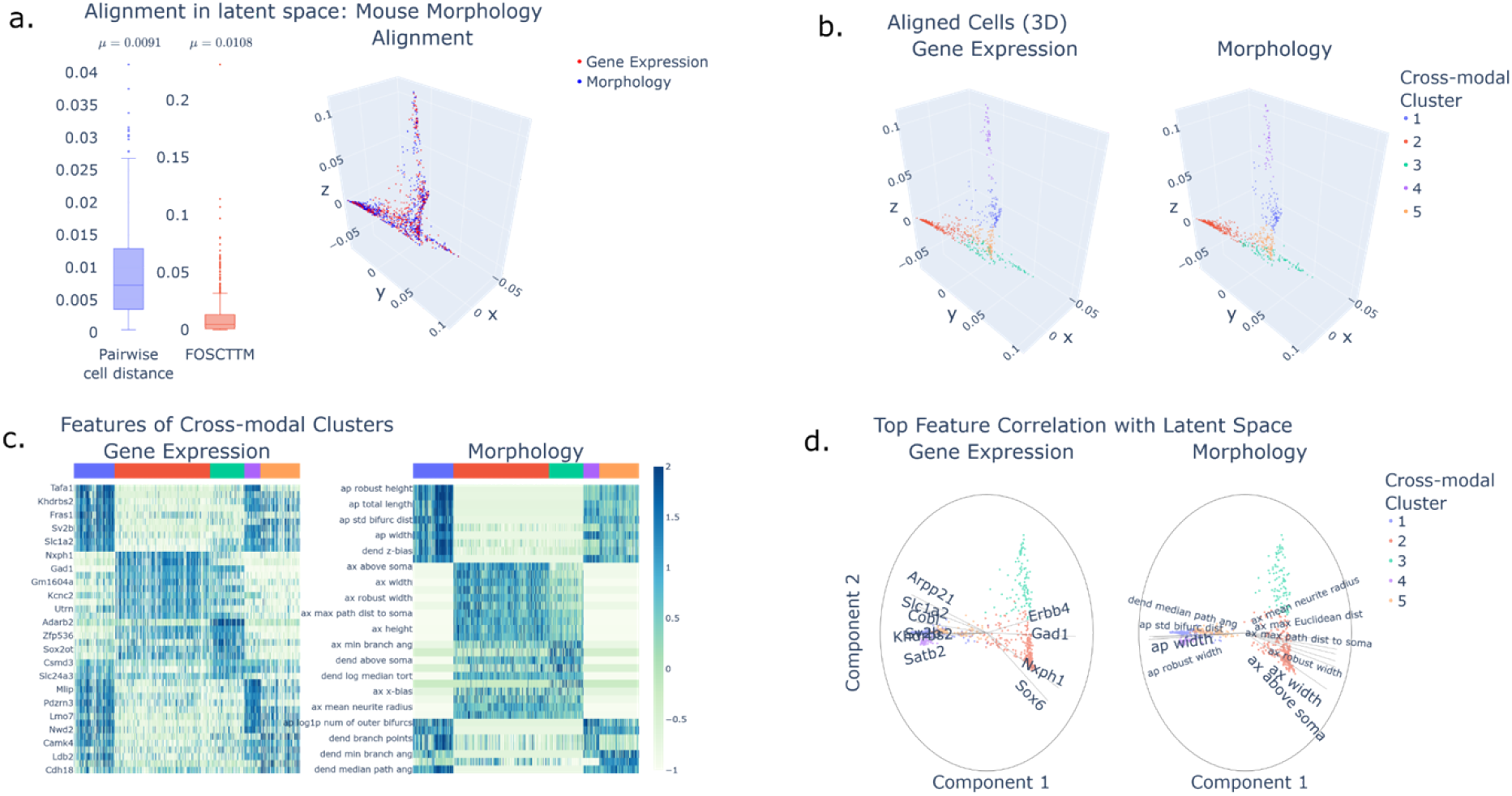
MANGEM analysis and visualization of neuronal gene expression and morphological features in mouse visual cortex. a.) Measures of alignment error and 3-d plot of superimposed aligned data in latent space are shown for the mouse morphology cortex dataset after nonlinear manifold alignment. b.) Cross-modal clusters, obtained by Gaussian mixture model, are indicated by color in plots of aligned data for each modality. c.) Feature expression levels across all cells for the top 10 differentially expressed features for each cross-modal cluster. Normalized feature expression was ranked using the Wilcox Rank Sum test. Cross-modal clusters are identified by the colored bar at the top of each plot. d.) Biplots for Gene Expression and Electrophysiological features using dimensions 1 and 2 of the latent space. The top 15 features by correlation with the latent space are shown plotted as radial lines where the length is the value of correlation (max value 1).

MANGEM is then used to separate the data into 5 clusters using a gaussian mixture model. The clusters closely align with true cell types (**Fig. 5b**). Then, differentially expressed features for each cluster may be downloaded and used for downstream analysis. The expressed genes can then be analyzed for importance in brain function.

MANGEM identifies Pvalb, Vip, Lamp5, and Sst among the top 2 most differentially expressed genes over the 5 cell clusters (**Fig. 5c**). These genes are commonly used to identify cell-type (26). So, MANGEM can be used to automatically perform cell-type clustering on multimodal datasets. In addition, MANGEM identifies Adarb2 as a differentially expressed gene. Adarb2 has been found to distinguish between two major branches of inhibitory neurons (27).

MANGEM also allows users to create Bibiplots to Visualize features important to the latent space (**Fig. 5d**). These features which are highly correlated with the latent space (e.g., SOX6 and sp_width) may then be the focus of future data exploration.

## Availability

MANGEM is freely available for use at https://ctc.waisman.wisc.edu/mangem. The source code for MANGEM is released under the MIT License and is available for download at https://github.com/daifengwanglab/mangem.

## Future Directions

MANGEM is a user-friendly web application designed primarily for biologists and neuroscientists. The app comes with pre-selected general-purpose hyperparameters that can be fine-tuned by users to suit their needs. With the rapid advancements in multimodal machine learning (28), MANGEM is constantly evolving to offer more advanced alignment options.

At present, MANGEM can only work with pre-processed electrophysiological and morphology features, but future versions may incorporate methods like deep neural networks to work with raw data (e.g., electrophysiological time-series data) or other types of data, such as genomics, epigenomics, or images. MANGEM is also capable of incorporating emerging machine learning approaches to infer missing modalities and cross-modal correspondence (29).

MANGEM uses cloud-based computing, which in the future will enable distributed training, making computation faster and providing a smoother experience for users. To further improve the efficiency of the app, MANGEM can be optimized for parallel processing, allowing it to take advantage of multiple processors and GPUs for faster computation. In addition to its alignment capabilities, MANGEM also enables collaborative work and data sharing. The app provides a centralized repository for storing and sharing aligned data, with built-in privacy and security measures to protect sensitive data.

## Supporting Information

Supplemental Data - tutorial video

Supplemental Materials - Supplemental Figures 1-2, Supplemental Table 1

## Author’s Contributions

D.W. conceived the study. D.W., R.O. and N.K. designed the methodology, performed analysis and visualization. R.O. implemented the software. D.W., R.O., and N.K. edited and wrote the manuscript. All authors read and approved the final manuscript.

## Acknowledgments

The authors wish to thank all members of the Wang lab for insightful discussions on the work.

